# A Biological Signature for the Inhibition of Outer Membrane Lipoprotein Biogenesis

**DOI:** 10.1101/2022.03.18.484967

**Authors:** Kelly M. Lehman, Hannah C. Smith, Marcin Grabowicz

## Abstract

The outer membrane (OM) of Gram-negative bacteria is an essential organelle that acts as a formidable barrier to antibiotics. Increasingly prevalent resistance to existing drugs has exacerbated the need for antibiotic discovery efforts targeting the OM. Acylated proteins, known as lipoproteins, are essential in every pathway needed to build the OM. The central role of OM lipoproteins makes their biogenesis a uniquely attractive therapeutic target, but it also complicates *in vivo* identification of on-pathway inhibitors, as inhibition of OM lipoprotein biogenesis broadly disrupts OM assembly. Here, we use genetics to probe the eight essential proteins involved in OM lipoprotein biogenesis. We define a biological signature consisting of three simple assays that can characteristically identify OM lipoprotein biogenesis defects *in vivo*. The few known chemical inhibitors of OM lipoprotein biogenesis conform to the biological signature. We also examine MAC13243, a proposed inhibitor of OM lipoprotein biogenesis, and find that it fails to conform to the biological signature. Indeed, we demonstrate that MAC13243 activity relies entirely on a target outside of the OM lipoprotein biogenesis pathway. Hence, our signature offers simple tools to easily assess whether antibiotic lead compounds target an essential pathway that is the hub of OM assembly.

**IMPORTANCE:** Gram-negative bacteria have an outer membrane, which acts as a protective barrier and excludes many antibiotics. The limited number of antibiotics active against Gram-negative bacteria, along with rising rates of antibiotic resistance, highlights the need for efficient antibiotic discovery efforts. Unfortunately, finding the target of lead compounds, especially ones targeting outer membrane construction, remains difficult. The hub of outer membrane construction is the lipoprotein biogenesis pathway. We show that defects in this pathway result in a signature cellular response that can be used to quickly and accurately validate pathway inhibitors. Indeed, we found that MAC13243, a compound previously proposed to target outer membrane lipoprotein biogenesis, does not fit the signature, and we show that it instead targets an entirely different cellular pathway. Our findings offer a streamlined approach to discovery and validation of lead antibiotics against a conserved and essential pathway in Gram-negative bacteria.

## INTRODUCTION

Since the advent of antibiotics, treatment of infection has been a race against time. Once antibiotics are introduced clinically, bacteria often quickly develop resistance. Antibiotic discovery efforts, with an emphasis on novel bacterial targets, are essential to the continuation of the current medical treatment model for curing infections. Resistance among Gram-negative pathogens is particularly concerning, as discovery of novel antibiotic classes targeting these bacteria has proved especially difficult (1). Gram-negative bacteria, such as *Escherichia coli*, have an outer membrane (OM) that acts as a selective permeability barrier against extracellular onslaughts, such as host immune factors and antibiotics (2). Thus, the OM is a prime antibiotic target, both because it is essential and because it is a protective barrier, leading many recent antibiotic discovery efforts to focus on OM biogenesis (3, 4).

The OM is an asymmetric lipid bilayer. The inner leaflet consists of phospholipids, while the outer leaflet primarily consists of lipopolysaccharide (LPS) (5). Construction of the OM requires specialized machinery, particularly because highly hydrophobic proteins and lipids must, somehow, cross an aqueous periplasm (Fig. 1) (6). Three machines are largely responsible for OM biogenesis: the lipopolysaccharide transport (Lpt) machine shuttles LPS to the OM (7), the β-barrel assembly machine (Bam) folds β-barrel proteins into the OM (8), and the localization of lipoprotein (Lol) pathway traffics lipoproteins to the OM (9). Notably, Bam, Lpt, and Lol require at least one essential OM lipoprotein component: BamD, LptE, and LolB, respectively (10–12). Thus, OM lipoprotein biogenesis, comprised of the lipoprotein maturation and trafficking pathways, is key to construction and integrity of the OM.

**Figure 1.**
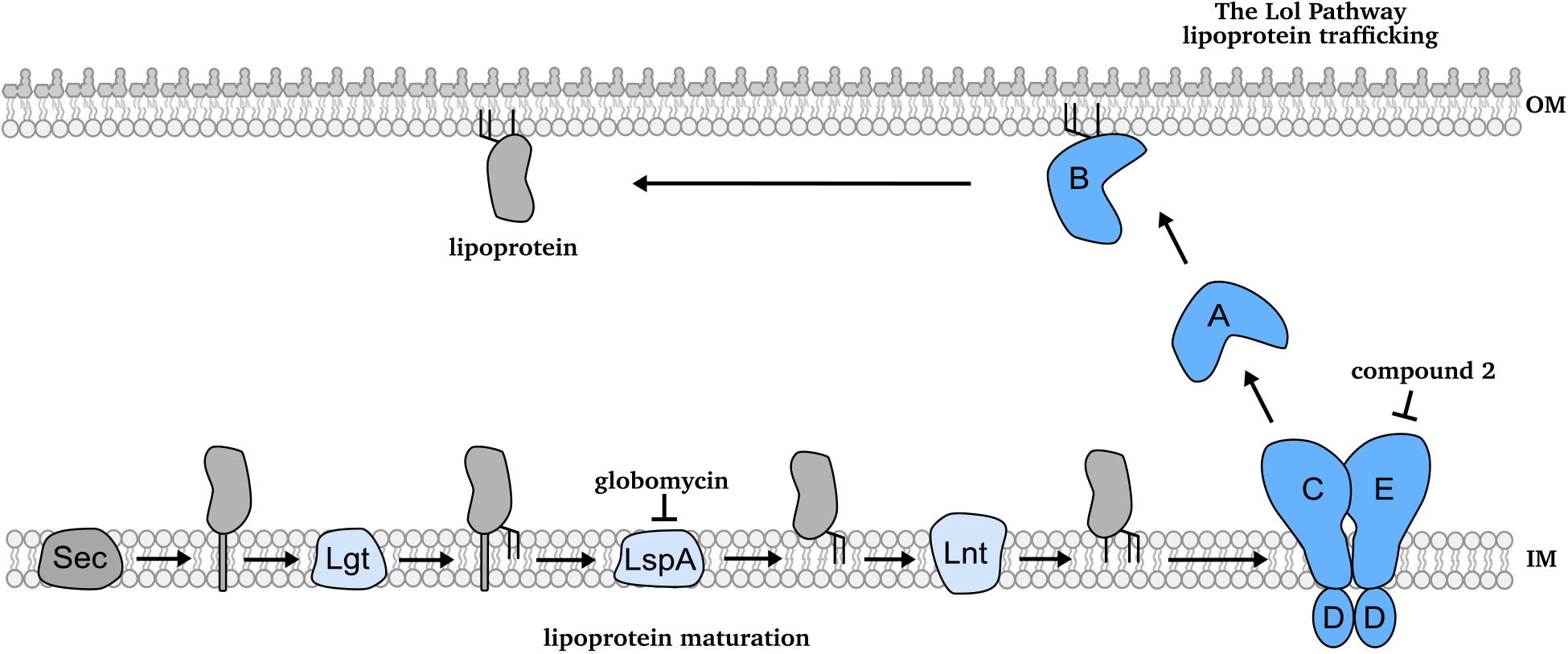
Lipoprotein maturation and trafficking and OM lipoprotein biogenesis inhibitors. Lipoproteins exit the cytoplasm via the Sec translocon, where they are tethered by their signal sequence in the inner membrane (IM). Before they are trafficked to the outer membrane (OM), lipoproteins must be modified by a series of lipoprotein maturation enzymes in the IM. Lipoproteins undergo sequential modifications by Lgt, LspA, and Lnt. Modified, triacylated lipoproteins are extracted by LolCDE. LolA receives lipoproteins from LolC, shielding their hydrophobic acyl chains as it traffics them across the aqueous periplasm. At the OM, LolB receives and inserts lipoproteins. Two known compounds inhibit lipoprotein maturation and trafficking: globomycin inhibits LspA, while compound 2 inhibits LolCDE.

All lipoproteins are synthesized in the cytoplasm and secreted. Lipoproteins destined for the OM must undergo a series of sequential modifications in the inner membrane (IM) before they are trafficked to the OM (Fig. 1) (13, 14). First, the enzyme Lgt transfers a diacylglycerol moiety from phosphatidylglycerol to an invariant cysteine of a target lipoprotein (15, 16). Next, the type II signal peptidase LspA cleaves the signal peptide (17). Finally, the acyltransferase Lnt adds a third acyl chain from phosphatidylethanolamine to the now N-terminal cysteine, producing a mature lipoprotein (18, 19). Lipoprotein maturation enzymes are highly conserved and essential among Gram-negative bacteria. However, some species can remain viable without *lnt* in laboratory conditions (20, 21).

A mature lipoprotein is trafficked to the OM if it contains residues specifying an OM localization signal, which varies across species (22–24). An ATP-binding cassette (ABC) transporter (LolCDE in *E. coli*) extracts mature, OM-targeted lipoproteins from the IM (25). Then, the chaperone LolA receives lipoproteins from LolC, shielding their hydrophobic acyl chains from the aqueous periplasm (26, 27). Finally, the OM lipoprotein LolB receives lipoproteins from LolA and inserts them into the OM (26). Many clinically important species produce LolB, although some Gram-negative species lack a clear homolog (9). A LolAB-independent trafficking mechanism also exists, though it alone cannot support viability in wildtype *E. coli* (28).

As OM assembly relies on lipoproteins, OM lipoprotein biogenesis is a crucial target for novel antibacterials. This pathway requires up to eight essential and conserved proteins, offering an array of potential therapeutic targets. In fact, a recent CRISPRi screen of the essential genes of *Vibrio cholerae* found that depletion of genes in the Lol trafficking pathway caused a more severe decrease in viability than any other essential genes (29).

Lipoproteins play an essential role in OM assembly, complicating unambiguous identification of OM lipoprotein biogenesis inhibitors *in vivo*. Inhibitors of OM lipoprotein biogenesis will wreak widespread havoc on β-barrel assembly, LPS transport, and cell wall biosynthesis. Lipoprotein trafficking and OM biogenesis are so entwined that lipoprotein trafficking inhibitors have emerged from screens designed to identify inhibitors of cell wall synthesis (30) and activators of s^E^, a monitor of β-barrel assembly (31).

*In vivo* target validation of new compounds active against essential pathways remains challenging. No protocol to validate inhibition of lipoprotein maturation or trafficking factors exists. In this work, we define a unique biological signature for target validation of OM lipoprotein biogenesis inhibitors in *E. coli*. Our signature consists of three biological effects that, collectively, are hallmarks of defective OM lipoprotein biogenesis: (i) increased OM permeability, (ii) toxicity of the major OM lipoprotein Lpp, and (iii) activation of the Cpx envelope stress response by a sensory OM lipoprotein, NlpE. We validate this signature using genetic depletions and chemical inhibitors (compound 2, globomycin) of essential steps in OM lipoprotein biogenesis. We then demonstrate the utility of our signature by examining MAC13243, a proposed LolA inhibitor, and find that MAC13243 fails to fulfill our biological signature. Finally, using genetics, we confirm that MAC13243 bioactivity is independent of LolA.

## RESULTS

### Depletion of OM lipoprotein biogenesis factors causes OM permeability

To establish a biological signature of lipoprotein maturation or trafficking inhibition, we used our current understanding of OM assembly to develop assays that report on OM lipoprotein biogenesis defects. Since at least one lipoprotein is essential to each OM biogenesis machine, we hypothesized that disrupting OM lipoprotein biogenesis would cause OM assembly defects that affect its antibiotic barrier function.

To assess OM barrier integrity when OM lipoprotein biogenesis is limited, we used a series of *E. coli* strains in which expression of OM lipoprotein biogenesis genes (*lspA, lolCDE, lolA, lolB*) depends on arabinose induction. Growth in media lacking arabinose depletes these essential proteins. We used checkerboard assays to measure sensitivity to three large scaffold antibiotics, which cannot pass through an intact OM, in response to depletion of OM lipoprotein biogenesis factors. Each antibiotic had a distinct target: novobiocin (a hydrophobic DNA gyrase inhibitor, Fig. 2), vancomycin (a hydrophilic cell wall biosynthesis inhibitor, Fig. 2), and rifampicin (a hydrophobic RNA polymerase inhibitor, Fig. S1).

**Figure 2.**
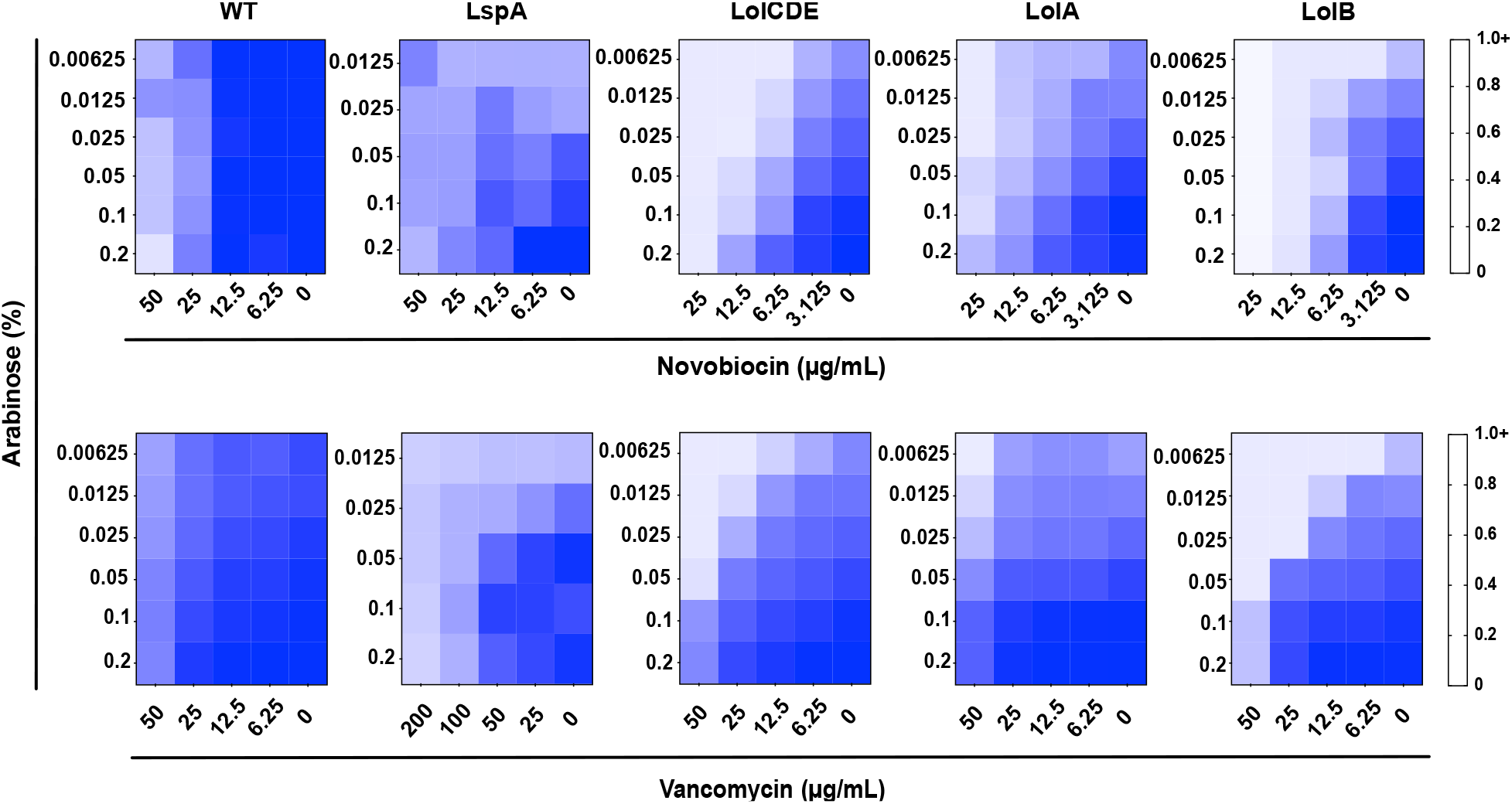
Depletion of lipoprotein maturation or trafficking factors causes outer membrane permeability. Strains in which LspA, LolCDE, LolA, or LolB were under an arabinose-dependent promoter were grown in decreasing concentrations of inducer and increasing concentrations of two large scaffold antibiotics, novobiocin and vancomycin. Depletion of any OM lipoprotein biogenesis factor tested caused increased sensitivity to large scaffold antibiotics. Arabinose did not affect the sensitivity of wildtype (WT) to large scaffold antibiotics. Data are from three independent experiments. Averaged density (A_600nm_) values of antibiotic-treated cultures relative to mock-treated control (set as 1.0) are shown.

As we depleted each protein, sensitivity to large scaffold antibiotics increased (Fig. 2 and Fig. S1). We observed variation in the extent of antibiotic sensitivity caused by depletion of each OM lipoprotein biogenesis factor, likely reflecting differing levels of depletion achievable with each construct. Nonetheless, depleting OM lipoprotein biogenesis increased OM permeability to antibiotics. The permeabilizing effect was compound-specific, as decreasing induction of *lspA, lolCDE, lolA*, or *lolB* did not sensitize cells to erythromycin (a hydrophobic macrolide inhibitor of translation, Fig. S1). Selective permeability caused by OM assembly mutants was previously observed and remains poorly understood (32). Our data confirm that defects in OM lipoprotein biogenesis weaken the integrity of the OM barrier.

### Loss of Lpp alleviates OM lipoprotein biogenesis defects

In addition to disrupting OM construction, OM lipoprotein biogenesis defects cause IM mislocalization of OM-targeted lipoproteins, which can be toxic (28). One such example is the OM lipoprotein Lpp, which covalently crosslinks to the cell wall from the OM, providing important architectural stability to the cell envelope (33–35). When Lpp is not trafficked efficiently, it accumulates in the IM and errantly crosslinks to peptidoglycan. Lpp crosslinking from the IM is lethal (36). Hence, although *lpp* is not essential, efficient OM lipoprotein biogenesis of Lpp is essential. We reasoned that Δ*lpp* would prevent toxicity, increasing viability when OM lipoprotein biogenesis is limited. We assessed the viability of Lgt, LspA, Lnt, LolCDE, LolA, or LolB depleted strains in the presence or absence of *lpp* using arabinose-inducible constructs (Fig. 3). Each gene is essential in both *lpp*^+^ and Δ*lpp* backgrounds; therefore, inducer-independent growth in these strains relies on leaky expression of the gene construct.

**Figure 3:**
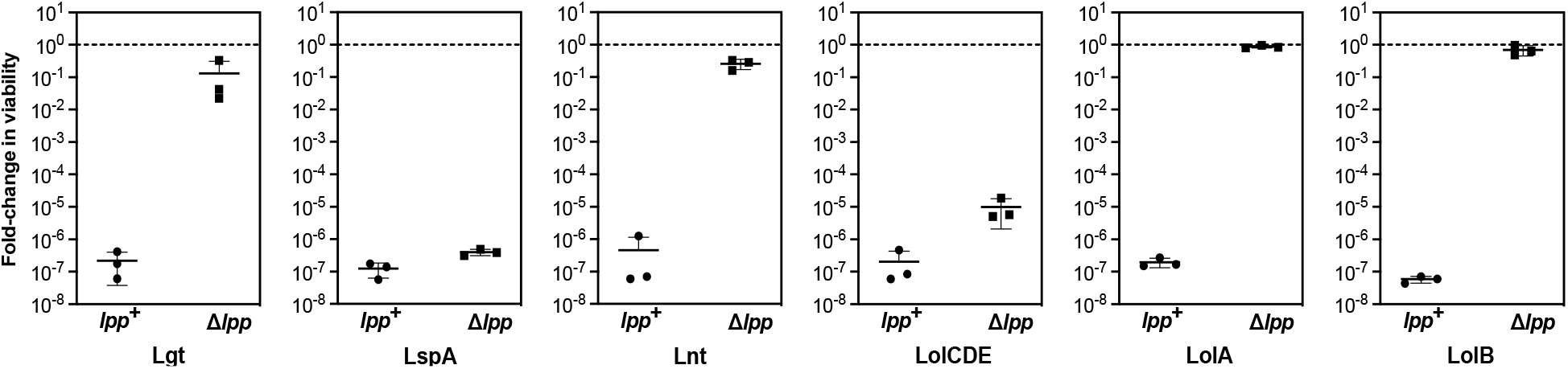
Deletion of *lpp* protects against lipoprotein maturation and trafficking defects. Relative viability of strains with arabinose-dependent expression of OM lipoprotein biogenesis proteins (Lgt, LspA, Lnt, LolCDE, LolA, or LolB). Viable counts per mL of culture were enumerated in the presence of 0.2% arabinose or in the absence of arabinose and used to determine the fold change in viability when inducer is absent. For LspA strains, the comparison was made between arabinose replete conditions (0.2%) and arabinose deplete (0.002%) conditions. Data represent three independent experiments and the mean.

Depletion of any OM lipoprotein biogenesis factor severely reduced viability of wildtype *E. coli*, as expected for essential genes. In all instances, Δ*lpp* improved viability without inducer. While Δ*lpp* caused striking increases in viability in Lgt-, Lnt-, LolA-, and LolB-depleted cells, we measured only modest increases in viability in LspA- and LolCDE-depleted cells. The variation in the alleviation of toxicity in Δ*lpp* strains likely reflects the dissimilar levels of depletion achievable with each gene construct, with little leaky expression of LspA or LolCDE. In fact, the tight regulation of the LspA depletion strain was previously demonstrated (37). Importantly, Δ*lpp* did not improve viability when essential components of the Bam and Lpt machines (BamD and LptE) were depleted (Fig. S2). Therefore, Δ*lpp* does not alleviate cell envelope defects in other essential OM assembly pathways. Rather, our data show that Δ*lpp* specifically improves the viability of cells when OM lipoprotein biogenesis is depleted.

### Depletion of OM lipoprotein biogenesis causes NlpE-dependent activation of Cpx

A series of stress responses monitor OM and cell envelope integrity (38). Among these is Cpx, a two-component system comprised of the histidine kinase CpxA and the response regulator CpxR (39). Together, CpxAR respond to OM perturbations and various other cellular signals (40). Cpx was recently shown to alleviate stress caused by defects in late steps of lipoprotein trafficking (28, 41, 42). We hypothesized that defective OM lipoprotein biogenesis would similarly activate Cpx, marking a signature of OM lipoprotein biogenesis stress.

To assess Cpx activation when OM lipoprotein biogenesis is defective, we used a reporter plasmid carrying a transcriptional *gfp* fusion to the promoter of the CpxAR-regulated gene *cpxP* (*P*_*cpxP*_*-gfp*). The plasmid was introduced into the LspA, LolCDE, LolA, and LolB depletion strains. We monitored GFP fluorescence as each OM lipoprotein biogenesis factor was depleted during sub-culture without inducer (Fig. 4). As expected, depletion of each OM lipoprotein biogenesis factor reduced growth. As growth slowed, we detected strong increases in fluorescence from *P*_*cpxP*_*-gfp* (Fig. 4), indicating activation of Cpx.

**Figure 4:**
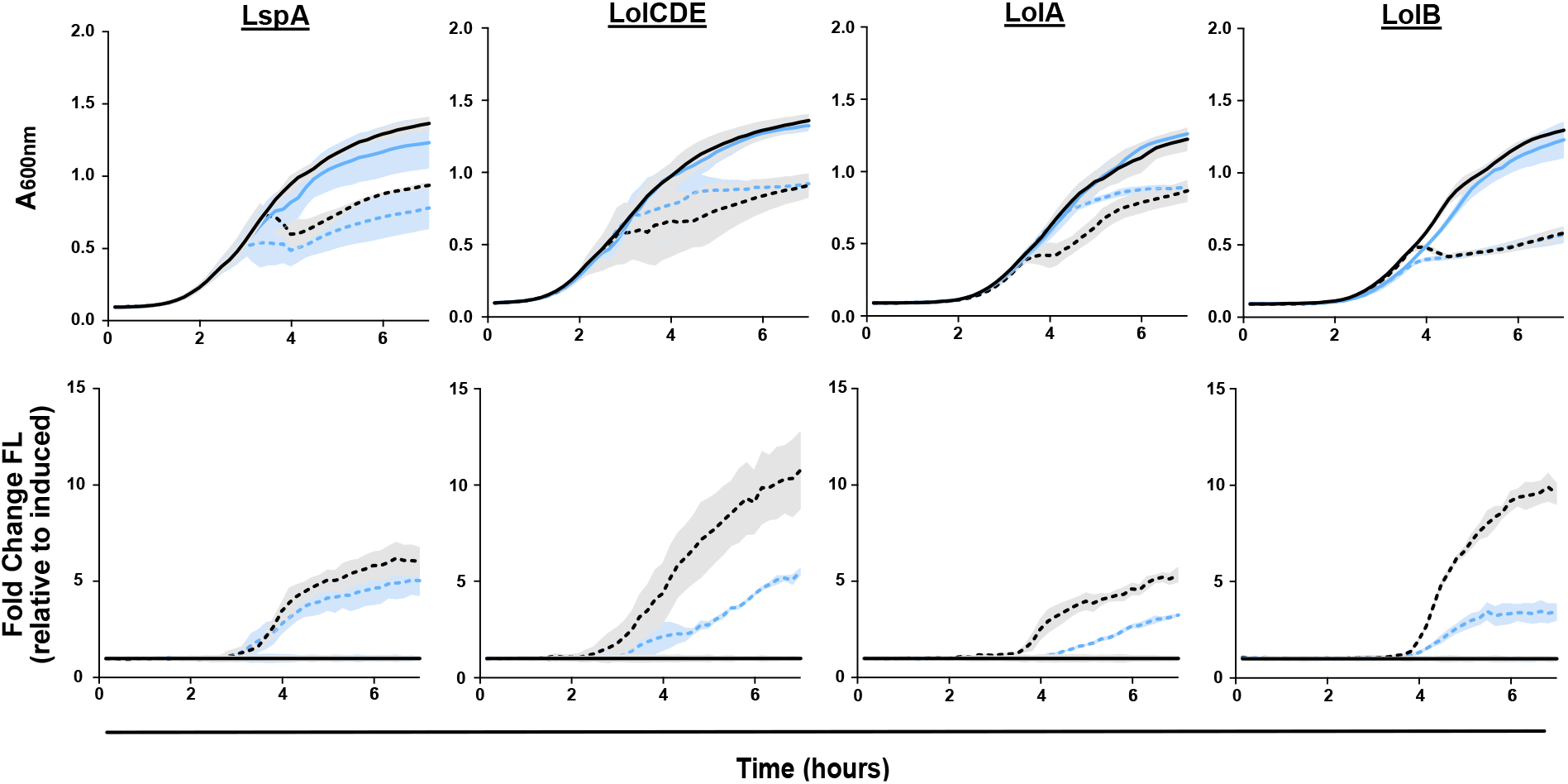
Depletion of lipoprotein maturation or trafficking causes NlpE-dependent activation of the Cpx stress response. Strains carrying *P*_*cpxP*_*-gfp* and inducible LspA, LolCDE, LolA, or LolB were grown with (solid) or without (dots) inducer (0.2% arabinose). Culture density (OD_600_, top) and fluorescence were measured to calculate fluorescence per cell (Fluorescence/OD_600_). Strains were tested in the presence (black) or absence (blue) of *nlpE*. (Data are average ± standard deviation, N=3)

As a variety of stimuli activates Cpx, we wanted to test whether the observed rapid and potent Cpx activation was specific to OM lipoprotein biogenesis defects. Recent work proposed that Cpx activation in response to defects in late OM lipoprotein biogenesis is due to mislocalization of the OM sensor lipoprotein NlpE to the IM (41, 42). We reasoned that if the observed Cpx activation was caused by sensing OM lipoprotein biogenesis defects, the early, strong Cpx activation would be NlpE-dependent. Hence, we deleted *nlpE* from our depletion strains and monitored expression from *P*_*cpxP*_*-gfp*. Growth of all strains was similar in the presence and absence of *nlpE*. Importantly, deletion of *nlpE* decreased fluorescence upon depletion of LolCDE, LolA, or LolB, indicating NlpE-dependent Cpx activation. We did not observe clear NlpE-dependent Cpx activation in the LspA strain, likely a product of the construct’s tight repression. However, as depletion of LolCDE, LolA, or LolB causes NlpE-dependent Cpx activation, we conclude that this is a strong indicator of OM lipoprotein biogenesis limitation.

### Stress responses with OM lipoprotein sensors are activated by limited OM lipoprotein biogenesis

Two other envelope stress responses monitor OM defects: Rcs and *σ*^E^. Rcs monitors defects through the OM lipoprotein RcsF (43), while *σ*^E^ directly detects misfolded β-barrel proteins. We reasoned that OM lipoprotein biogenesis defects would lead to early activation of Rcs but would not activate *σ*^E^, as no lipoprotein is involved in the *σ*^E^ response. To assess Rcs and *σ*^E^ activation, we introduced reporter plasmids carrying a transcriptional *gfp* fusion to the Rcs-responsive *osmB* promoter (*P*_*osmB*_*-gfp*) or the *σ*^E^-dependent *micA* promoter (*P*_*micA*_*-gfp*) (Fig. S3) (44, 45). We also constructed a control plasmid expressing GFP from a housekeeping RpoD-dependent promoter (*P*_*rpoD*_*-gfp*) to control for artifactual increases in fluorescence (Fig. S3). We introduced each plasmid into the LspA, LolCDE, LolA, and LolB depletion strains and measured growth and GFP fluorescence during depletion. We observed increases in fluorescence from *P*_*osmB*_*-gfp* when OM lipoprotein biogenesis factors were depleted, indicating Rcs activation. Conversely, depletion did not strongly activate *P*_*micA*_*-gfp*, with the exception of LspA. None of the depletion strains caused strong activation of the control reporter, *P*_*rpoD*_*-gfp*. Thus, we propose that the specific activation of Cpx and Rcs is a strong indicator of OM lipoprotein biogenesis inhibition, while an absence of *σ*^E^ activation is important for discrimination between specific OM lipoprotein defects and generalized cell envelope defects.

### Chemical inhibitors of OM lipoprotein biogenesis conform to the biological signature

Genetic depletions allowed us to establish three signature hallmarks of defects in OM lipoprotein biogenesis: (i) OM permeabilization, (ii) Lpp toxicity, (iii) and NlpE-specific activation of Cpx. We next tested our signature using chemical inhibition of the OM lipoprotein biogenesis pathway. We used two well-characterized compounds: globomycin (46) and the pyridine-imidazole “compound 2” (47) (Fig. 1). Globomycin inhibits LspA (48), while compound 2 inhibits LolCDE function (47).

To probe OM permeability, we assessed sensitivity to large scaffold antibiotics upon treatment with globomycin or compound 2 using checkerboard assays. As globomycin and compound 2 poorly penetrate *E. coli*, we tested a Δ*tolC* strain in which antibiotic efflux is inactivated. A Δ*tolC* strain should allow cellular accumulation of both compounds. Treatment with globomycin or compound 2 sensitized cells to vancomycin, indicating decreased integrity of the OM barrier (Fig. 5). Thus, both compounds satisfied the first criterium of the proposed biological signature.

**Figure 5:**
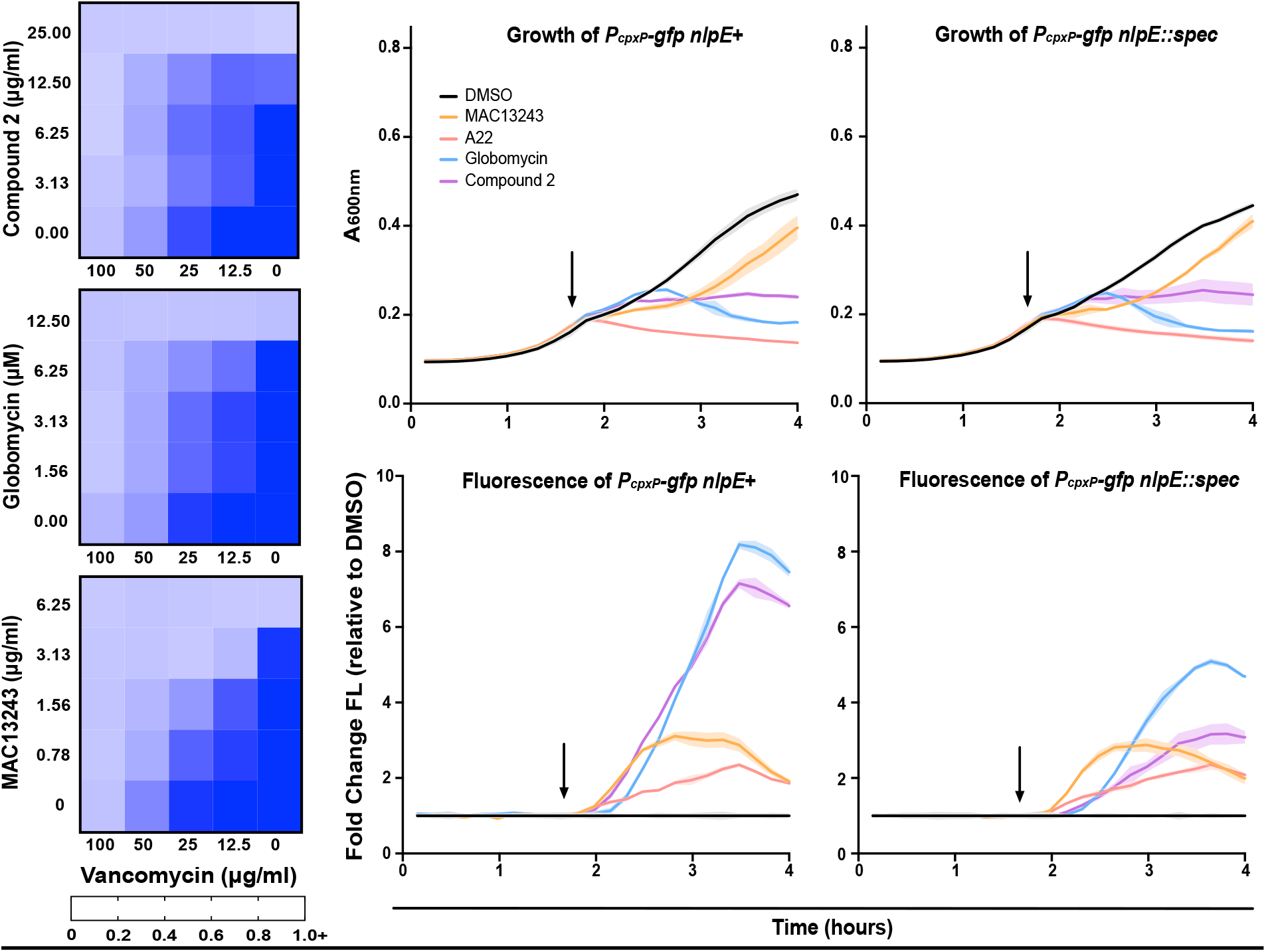
Chemical inhibitors of lipoprotein maturation or trafficking fit the expected profile of OM lipoprotein biogenesis inhibition. (Left) OM permeability was assessed in efflux defective mutants (Δ*tolC*) by treatment with increasing concentrations of vancomycin and compound 2 (top), globomycin (middle), or MAC13243 (bottom). Data are from three independent experiments. Averaged density (A_600nm_) values of antibiotic-treated cultures relative to mock-treated control (set as 1.0) are shown. (Right) Culture density (OD_600_, top) and fluorescence were used to calculate fluorescence per cell in Δ*tolC* strains with native *nlpE* or with a chromosomal deletion of *nlpE* (*nlpE::spec*). Cells were treated after 100 minutes (arrow), with DMSO (black), MAC13243 (orange), A22 (red), globomycin (blue), or compound 2 (purple). (Data are average ± standard deviation, N=3)

We next tested whether Δ*lpp* was protective against treatment with either inhibitor. In agreement with previous work, Δ*lpp* increased the MIC of globomycin and compound 2 (Table 1) (47, 49). Therefore, both compounds fulfill the second criterium of our signature.

**Table 1:** Deletion of *lpp* increases resistance to OM lipoprotein biogenesis inhibitors.

| Strain | Globomycin (μM) | Compound 2 (μg/mL) | A22 (μg/mL) | MAC13243 (μg/mL) |
| --- | --- | --- | --- | --- |
| *ΔtolC lpp+* | 12.5 | 10 | 2.5 | 2.5 |
| *ΔtolC Δlpp* | >50 | >40 | 2.5 | 2.5 |

Finally, we tested stress response activation upon treatment with both inhibitors. Strains were treated with globomycin or compound 2 after 100 minutes of growth. Both compounds inhibited growth similarly. Almost immediately after treatment with either compound, we detected a strong increase in fluorescence from a P_*cpxP*_*-gfp* reporter (Fig 5). Deletion of *nlpE* delayed and strongly reduced GFP fluorescence upon treatment with either compound. We found that globomycin induced fluorescence from Rcs-activated *P*_*osmB*_*-gfp*, consistent with prior observations (Fig. S4). Treatment with compound 2 also increased fluorescence of *P*_*osmB*_*-gfp* (Fig. S4). However, both globomycin and compound 2 caused little activation of the *σ*^E^-dependent *micA* promoter (*P*_*micA*_*-gfp*) (Fig. S4). Thus, both globomycin and compound 2 satisfy all three of our criteria and fully conform to our biological signature. Collectively, our data demonstrate that OM lipoprotein biogenesis defects, whether induced genetically or with chemical inhibitors, produce a distinctive biological signature of OM lipoprotein biogenesis inhibition.

### Proposed LolA inhibitor MAC13243 does not fit the biological signature

Recent work proposed that MAC13243 inhibits LolA (50). Indeed, MAC13243 permeabilizes *E. coli* to large scaffold antibiotics (51), and LolA over-production protects against MAC13243 (50). Curiously, MAC13243 degrades into a thiourea compound closely related to A22, an inhibitor of the essential cytoskeletal protein MreB (52–54). *In vitro*, MAC13243 and A22 interact with purified LolA (54). However, clear *in vivo* LolA inhibition has yet to be demonstrated. We hypothesized that, if they target LolA *in vivo*, both MAC13243 and A22 would fit our biological signature.

As a control for LolA inhibition, we also designed an allele-specific system for inhibiting LolA. First, we introduced a V24C substitution in LolA. The V24 residue is proposed to be important to lipoprotein binding by LolA (26). A plasmid carrying *lolA(V24C)* complemented deletion of native *lolA*, indicating that the mutation does not reduce LolA activity. To inhibit LolA(V24C), we treated cells with the thiol-reactive compound 2-[(methylsulfonyl)thio]-ethanesulfonic acid (MTSES). Previous work illustrated that, despite the potential effect of MTSES on any thiol group in the cell, clever introduction of cysteines at key sites causes protein-specific sensitivity to MTSES (55, 56). We reasoned that V24C would introduce an MTSES target within a functionally important region of LolA. Indeed, *lolA*(V24C) was more sensitive to MTSES than the *lolA*^*+*^ (Table S1). Treatment with MTSES caused only minor growth defects in the *lolA*^*+*^ strain, yet the same treatment was lethal in the *lolA(V24C)* strain (Fig. S5). Hence, MTSES allowed us to semi-selectively inhibit LolA in strains producing the V24C variant.

We tested MAC13243, A22, and our allele-specific MTSES inhibitor system using our biological signature. Treatment with MAC13243 increased sensitivity to novobiocin, vancomycin, and rifampicin, indicating increased OM permeability (Fig. 5). This is in keeping with previous observations that MAC13243 permeabilizes *E. coli* to vancomycin (51) and observations that A22 permeabilizes *E. coli* to novobiocin (57). Increasing concentrations of MTSES also increased sensitivity of LolA(V24C) to large scaffold antibiotics (Fig. 6, Fig. S6).

**Figure 6:**
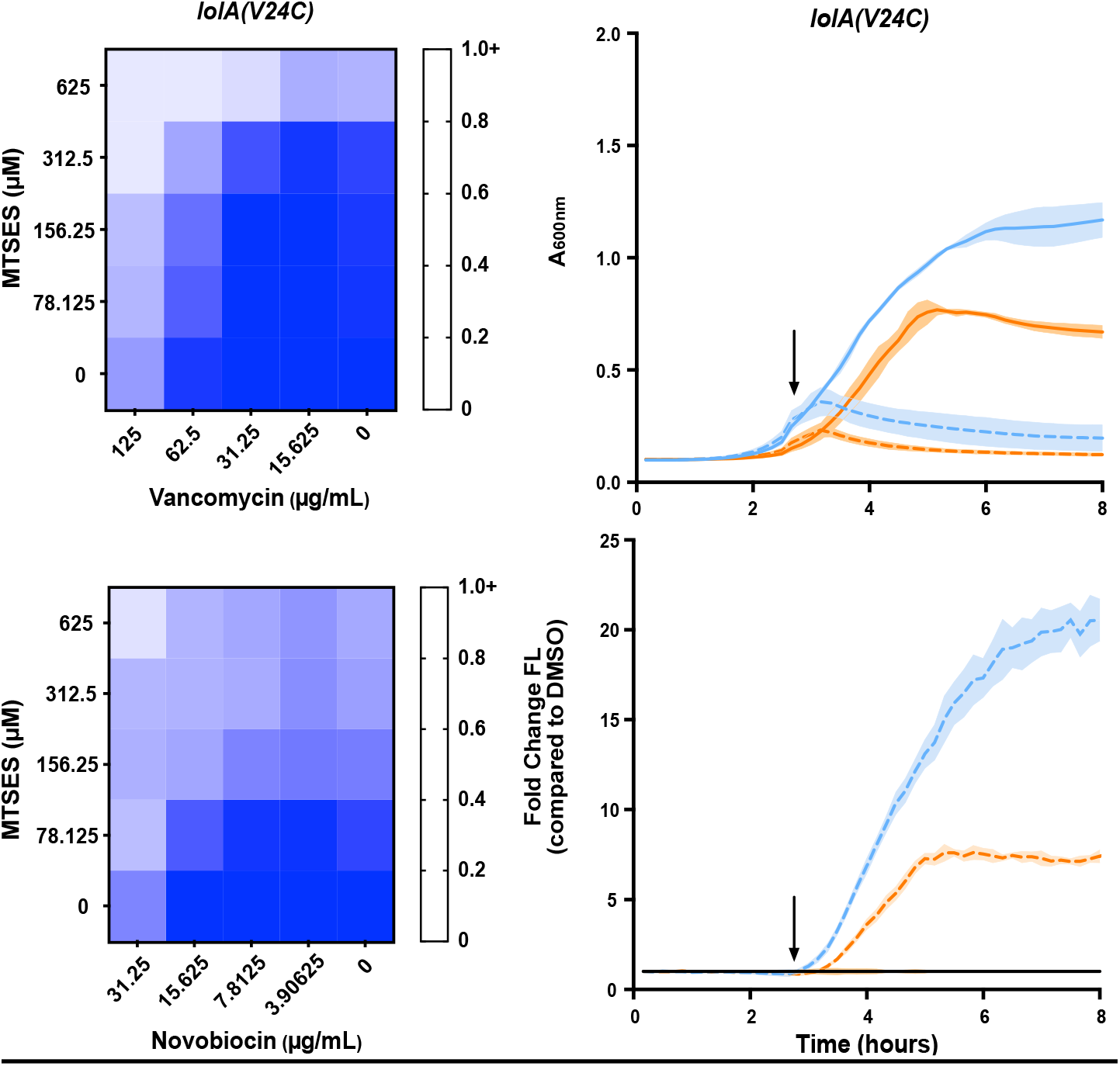
An allele specific inhibitor of LolA causes OM permeability and activation of the Cpx stress response. **(A)** OM permeability to vancomycin and novobiocin was assessed in a strain carrying *lolA(V24C)* upon treatment with increasing concentrations of MTSES. Data are from three independent experiments. Averaged density (A_600nm_) values of antibiotic-treated cultures relative to mock-treated control (set as 1.0) are shown. (B) Growth (OD_600_) of strains carrying *lolA(V24C)* and a Cpx reporter plasmid (*P*_*cpxP*_*-gfp*) was measured. Strains either had native *nlpE* (blue) or were Δ*nlpE::spec* (orange). In early log phase (arrow), strains were treated with 0.5 mM MTSES (dotted) or vehicle control (1% DMSO) (solid). Fluorescence was measured and normalized to OD_600_ to calculate fluorescence per cell. Reporter values were normalized to a DMSO-treated control (data are average ± standard deviation, N=3).

Since Δ*lpp* vastly improves viability when LolA is depleted, we expected Δ*lpp* would make *E. coli* more tolerant to a compound targeting LolA. Indeed, Δ*lpp* increased the MIC of MTSES in the *lolA(V24C)* strain (Table S1). However, Δ*lpp* had no effect on the MIC of MAC13243 or A22 (Table 1). This suggests that toxic mislocalization of Lpp is not a significant contributor to the lethality of MAC13243 or A22.

Next, we evaluated Cpx activation upon chemical inhibition of LolA. Treatment of *lolA(V24C)* with MTSES caused rapid growth arrest and strong activation of Cpx, just as we observed upon LolA depletion. Moreover, this activation was clearly NlpE-dependent (Fig. 6, Fig. S7). Treatment with MAC13243 or A22 also caused rapid growth inhibition and strong Cpx activation (Fig. 5). However, Cpx activation in response to MAC13243 or A22 treatment was entirely NlpE-independent (Fig. 5). Analysis of Rcs activation showed that MTSES induced Rcs in a non allele-specific manner (Fig. S8). Both MAC13243 and A22 caused delayed Rcs activation (Fig. S4).

Thus, MAC13243 and A22 fail to meet the biological signature of OM lipoprotein biogenesis inhibition. Deletion of *lpp* does not alleviate lethal effects of either compound, and both compounds activate Cpx in an NlpE-independent manner. Our data suggest that treatment with these compounds does not appreciably inhibit OM lipoprotein biogenesis, suggesting LolA is not inhibited *in vivo*.

### MAC13243 activity is LolA-independent

We sought to conclusively assess if the biological activity of MAC13243 occurs through inhibition of LolA *in vivo*. While *lolA* is essential in wildtype *E. coli*, genetic conditions exist under which both *lolA* and *lolB* can be deleted (28). As LolA and LolB work in concert, we examined the activity of MAC13243 in a strain lacking both LolA and LolB (Δ*lolAB*). We expected that MAC13243 would affect cells that produce LolA and LolB (*lolAB*^+^) but would not show activity in cells that lack the proposed LolA target (Δ*lolAB*). Surprisingly, we observed that MAC13243 causes OM permeabilization even in the absence of LolA (Fig. 7). Thus, the OM permeabilizing effect of MAC13243 is not dependent on LolA inhibition.

**Figure 7:**
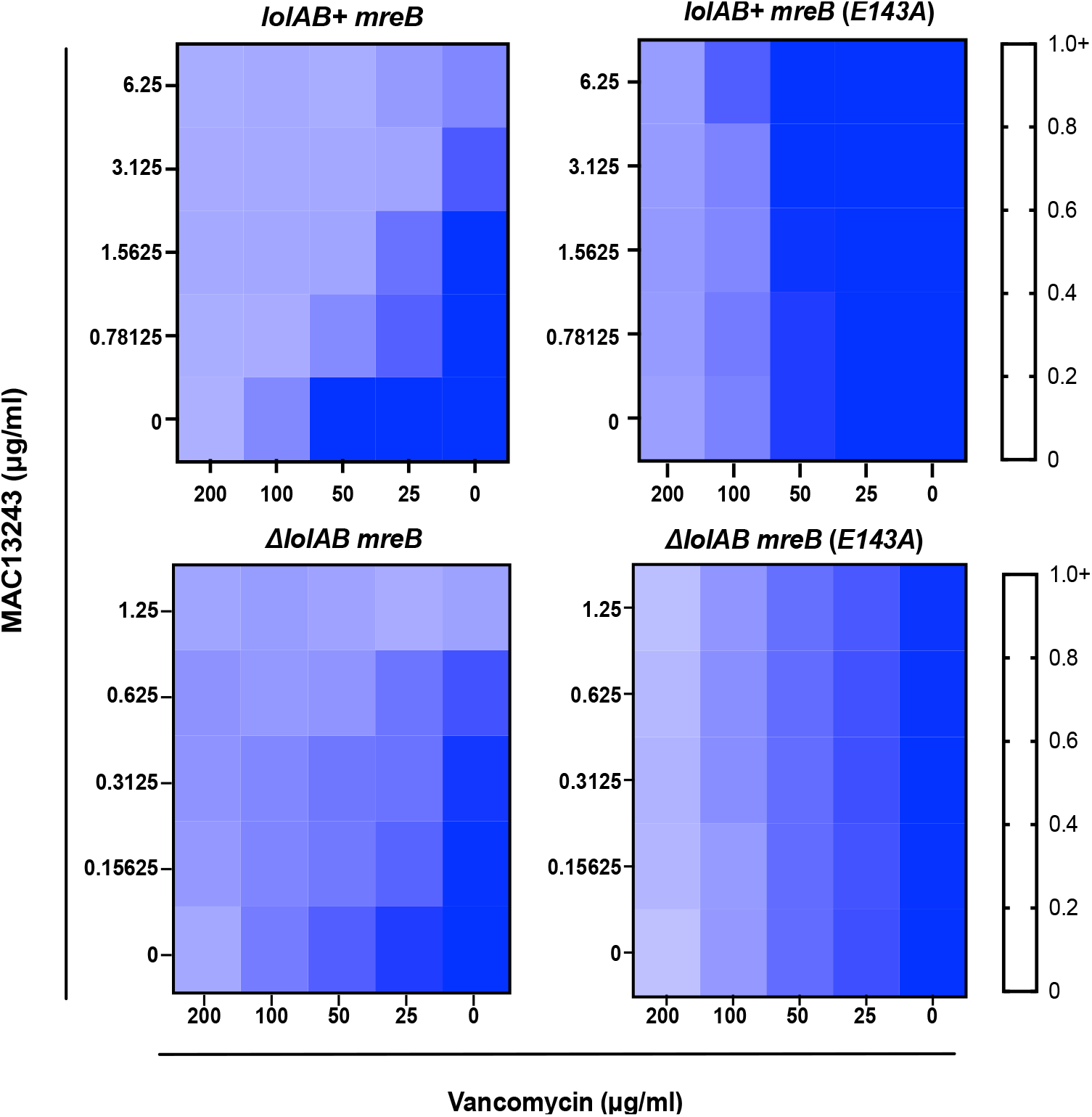
MAC13243 activity is independent of LolA. To test MAC13243 activity on LolA, OM permeability of strains in which chromosomal *lolAB* (*lolAB+*) are present or absent (Δ*lolAB*) was assessed. Strains carried either wildtype *mreB* or *mreB(E143A)* and were assessed upon treatment with increasing concentrations of MAC13243 and vancomycin. Data are from three independent experiments. Averaged density (A_600nm_) values of antibiotic-treated cultures relative to mock-treated control (set as 1.0) are shown.

The inhibition of MreB by A22 is well characterized. Since MAC13243 is chemically similar to A22, we examined whether the permeabilizing effect of MAC13243 relied on MreB inhibition. An E143A substitution in MreB confers resistance to A22, likely by preventing its binding (58). Interestingly, an *mreB(E143A)* allele also increased resistance to MAC13243. Moreover, in a *mreB(E143A)* background, MAC13243 did not permeabilize the OM to large scaffold antibiotics (Fig. 7). Hence, the activity of MAC13243 was entirely dependent on a susceptible MreB protein and independent of the presence of LolA in the cell. Collectively, our data strongly argue that the *in vivo* target of MAC13243 is MreB, not LolA.

## DISCUSSION

OM lipoprotein biogenesis is an attractive antibiotic target, as it is required for OM construction and integrity. However, there is currently no protocol for validating lipoprotein maturation or trafficking inhibitors. Herein, we establish a three-fold biological signature of OM lipoprotein biogenesis limitation: (i) permeabilization of the OM to large scaffold antibiotics, (ii) toxicity of Lpp, and (iii) NlpE-dependent activation of Cpx. This signature can be used to validate OM lipoprotein biogenesis inhibitors *in vivo*. Indeed, known inhibitors fully conform to this signature.

The first parameter of our biological signature is OM permeabilization. Prior work firmly established that mutations in the Bam and Lpt machines permeabilize the OM to antibiotics (59). This property has been exploited for genetic analysis of Bam and Lpt. Our data now show that the same chemical genetic logic extends to the OM lipoprotein biogenesis. Increased OM permeability is arguably the least discerning parameter in our biological signature, since permeability can be expected in response to defects in OM assembly, cell wall synthesis, or antibiotic efflux. As such, we see OM permeability as a primary classifier, which, if not satisfied, can exclude compounds that do not target OM lipoprotein biogenesis.

The second parameter of our biological signature relies on increased viability in the absence of Lpp. Notably, we show that Δ*lpp* alleviates defects in any stage of OM lipoprotein biogenesis yet does not alleviate defects in other OM assembly pathways (Bam and Lpt). The covalent linkage between OM-localized Lpp and cell wall peptidoglycan serves an important role in cell envelope architecture (34, 35). However, when Lpp cross-links from the IM, it is lethal to the cell (36). Defects at any stage in OM lipoprotein biogenesis should cause Lpp to accumulate in the IM, and deletion of *lpp* prevents lethal toxicity. In fact, *lpp* mutations alleviate temperature-sensitivity of *E. coli* or *Salmonella lgt* mutations (60, 61) and confer resistance to globomycin or LolCDE-targeting chemical inhibitors (47, 49).

Loss-of-function *lpp* mutations can be isolated with high frequency in the laboratory, suggesting a ready genetic route for resistance to novel therapeutics targeting OM lipoprotein biogenesis. Yet, it is unclear that similar *lpp* mutations could be isolated in a clinical context. The absence of Lpp dysregulates the cell envelope architecture, which leads to excessive OM blebbing and hypersensitivity to detergents frequently encountered by enteric bacteria, such as bile salts (62). Indeed, Δ*lpp* mutants survive poorly in mammalian hosts and are highly sensitive to complement-mediated immune clearance in serum (63–67). Therefore, although *lpp* is not essential in the laboratory, there is strong evidence to suggest that *lpp* is essential for infection. It is highly unlikely that *lpp* mutations could arise inside patients or animals treated with OM lipoprotein biogenesis inhibitors. Similarly, *Acinetobacter baumannii* mutants that no longer produce lipooligosaccharide can be readily isolated in the lab following colistin selection, but no such mutants have been recovered clinically from colistin-treated patients (68, 69).

A recent study described a macrocyclic peptide (G2824) that inhibits Lgt activity and is bactericidal to *E. coli* (70). Notably, Δ*lpp* did not confer resistance to G2824. In fact, Δ*lpp* sensitized bacteria to G2824. This is unexpected in light of our data showing that Δ*lpp* significantly increased viability of Lgt-depleted *E. coli* and other studies reporting that *lpp* mutations alleviate the effects of defective *lgt* alleles (60, 61). G2824 has two reported activities: it impedes lipoprotein modification by Lgt, and it prevents Lpp attachment to peptidoglycan. This dual activity suggests G2824 may have multiple targets *in vivo*. Both of the inhibited reactions, Lgt modification and Lpp attachment to peptidoglycan by the L,D-transpeptidases LdtABC, rely on cysteine residues. If G2824 has affinity for cysteines in the periplasm, it would interfere with both lipoprotein maturation and Lpp-peptidoglycan attachment, as reported. This hypothesis requires testing, but such a generalized activity of G2824 in the cell envelope would explain why Δ*lpp* sensitizes *E. coli* treated with G2824. The absence of Lpp causes severe envelope disruption that is exacerbated by inhibiting transpeptidases and cysteine-dependent periplasmic reactions (71).

The final parameter of our biological signature of OM lipoprotein biogenesis inhibition is NlpE-dependent activation of Cpx. Recent work revealed that NlpE acts as a real-time sensor of lipoprotein stress (41, 72). When lipoprotein trafficking is disrupted, NlpE becomes trapped in the IM, where it signals to CpxA. In keeping with this model, we found that depletion or chemical inhibition of OM lipoprotein biogenesis causes NlpE-dependent activation of Cpx. As lipoprotein trafficking is just one of the stressors to which CpxAR responds, general cell envelope defects likely still activate Cpx, yet they do so independently of NlpE. Although LspA depletion only caused NlpE-independent Cpx activation, globomycin, a well-studied LspA inhibitor, caused clear NlpE-dependent Cpx activation. The conformity of globomycin to our biological signature indicates the usefulness of our assay for validation of OM lipoprotein biogenesis inhibitors. NlpE allows rapid, robust activation of Cpx, speaking to its imperative role reacting to OM lipoprotein biogenesis stress and its usefulness as a criterium in the biological signature of OM lipoprotein biogenesis inhibition.

In addition to Cpx activation, our data indicate that Rcs activation is a strong indicator of OM lipoprotein biogenesis inhibition. Stress response activation is, thus, a powerful tool for the identification and validation of OM biogenesis inhibition. However, as measuring the NlpE-dependence of Cpx activation provides direct assessment of OM lipoprotein biogenesis, we found it to be the most informative parameter for identification of OM lipoprotein biogenesis inhibitors.

Finally, our results offer an essential conclusion to an ongoing discussion of the true target of MAC13243 *in vivo*. MAC13243 was originally discovered using overexpression of the essential genes of *E. coli* (50). Overexpression of LolA protected against treatment with MAC13243 (50). Later studies found that MAC13243 degrades under aqueous conditions into S-(4-chlorobenzyl)isothiourea, a close analog of the known MreB inhibitor A22 (54). *In vitro*, MAC13243, its S-(4-chlorobenzyl)isothiourea derivative, and A22 were all suggested to bind purified LolA (54). Given this evidence, MAC13243 has been embraced in the field as a LolA inhibitor (73–76). Our results, however, indicate that neither MAC13243 nor A22 conform to the expected signature of an OM lipoprotein biogenesis inhibitor. As Δ*lpp* offers no protection and Cpx activation is NlpE-independent in response to MAC13243 or A22 treatment, we suggest that neither compound appreciably impedes LolA activity *in vivo*. MAC13243 and A22 only conform to one criterium of our signature: OM permeabilization. Interestingly, we found that MAC13243 still causes OM permeabilization in the absence of LolA. Conversely, OM permeabilization does require a susceptible allele of *mreB*.

Comparing otherwise isogenic *lolAB*^*+*^ and Δ*lolAB* strains, we detected an increase in sensitivity to vancomycin. Hence, the loss of the LolAB trafficking pathway caused additional antibiotic sensitivity. We would expect that a compound that inhibits LolA should similarly sensitize to vancomycin. However, we failed to see any sensitization to vancomycin, even at high concentrations of MAC13243, in either *mreB*^+^ or *mreB(E143A)*. Collectively our data strongly argue against any *in vivo* activity of MAC13243 against LolA. Recent evidence also supports this conclusion. Overexpression of an inhibitor’s target can confer resistance to some inhibitors. This was the interpretation originally used to explain how LolA overexpression provides resistance to MAC13243. However, recent work found that LolA overexpression triggers activation of Rcs, explaining the protective effect of LolA overexpression again MAC13243 (77). Inactivation of Rcs abolished the protective effect of LolA overexpression. Thus, it is Rcs activation, not LolA overexpression, that is protective. Given our evidence, MAC13243 should be classified as an MreB-inhibiting compound, since it has no apparent activity against LolA *in vivo*.

We propose that the described biological signature discerns between those compounds that specifically inhibit OM lipoprotein biogenesis and those that interfere with the closely related processes of OM or cell envelope assembly. In an age of increasing antibiotic resistance, discovery efforts are imperative, yet lead compound target validation *in vivo*, especially for compounds targeting essential proteins and processes, remains challenging. Several groups have recently established clever methods to act as roadmaps for discovery and validation of on-pathway inhibitors of a variety of cellular pathways, including β-barrel assembly and cell elongation (77, 78). Our biological signature of OM lipoprotein biogenesis adds to this suite of resources, providing an invaluable tool for rapid validation of inhibitors of OM lipoprotein biogenesis.

## MATERIALS AND METHODS

### Strain Construction

Strains and plasmids used are provided in Tables S2 and S3, respectively. Oligonucleotides used in constructing strains and plasmids are provided in Table S4. Strains were grown in Lennox Broth (LB) supplemented with ampicillin (Amp, 125 mg/L), spectinomycin (Spec, 50 mg/L), kanamycin (Kan, 25 mg/L) or arabinose (Ara, 0.2% w/v) as needed. The *tolC* and *lpp* kanamycin-resistant deletion-insertion mutants were obtained from the Keio collection (79). The Δ*lolA*::*kan*, Δ*lolB*::*kan*, Δ*nlpE*::*spec*, and Δ*lolCDE*::*cam* alleles were previously described (28, 80). Deletion-insertion mutations and the complementing constructs of *lspA, lnt*, and *lgt* have also been previously described (37, 81, 82). A22-resistant *mreB(E143A)* was previously described (58). Strains were constructed by standard P1*vir* transduction of antibiotic resistance-marked alleles or by standard plasmid transformations.

### Checkerboard Assays

Overnight cultures were diluted to OD_600_=0.1, then further diluted 1:1,000 into fresh broth. For LspA, LolCDE, LolA, and LolB depletion strains, 60 μL of subculture was added to each well of a 96-well microtiter plate. Next, varying amounts of arabinose (diluted in LB broth) were added in a volume of 20 μL. Finally, varying concentrations of antibiotic (diluted in LB broth) were added in a volume of 20 μL. Plates were sealed with Breathe-Easy Gas Permeable Film (Sigma Z380059) and incubated overnight at 37°C. For checkerboard assays using MTSES, subcultures were prepared as above in 60 μL LB with 0.2% arabinose. To each well, varying MTSES amounts (20 μL) and varying antibiotic amounts (20 μL) were added. Plates were incubated for 48 hours at 30°C. For MAC13243 checkerboard assays, cultures were prepared as above in 60 μL volume (without arabinose). Varying amounts of MAC13243 (in 20 μL) and antibiotic (in 20 μL) were added to each well. Checkerboard assays using globomycin and compound 2 were prepared as above but scaled down to 40 μL final volume in a 384-well microtiter plate. In all cases, A_600nm_ was read using a Synergy H1 microplate reader (Biotek).

### MTSES growth curves and GFP reporter assays

Overnight cultures were diluted 1:1,000 into LB broth supplemented with arabinose (0.02%) and Kan, where appropriate. Aliquots of 1.96 mL seeded each well of a 24-well microtiter plate. Plates were sealed with breathable film and incubated at 37°C with shaking in a Synergy H1, measuring A_600nm_. After 180 mins, MTSES or vehicle control DMSO was added in a volume of 40 μl. Plates were then returned to the plate reader.

### Cell Viability Assays

Overnight cultures of depletion strains (*lpp*^+^ or Δ*lpp*) were grown in LB supplemented with 0.2% arabinose. Dilutions of the saturated culture were plated onto LB with arabinose and LB alone. Plates were incubated at 37°C overnight. Viable counts were enumerated as colony-forming units per mL of culture. The ratio of viable cells in the presence or absence of arabinose was calculated.

### MIC assays

Cultures (5 × 10^4^ cells/mL) were seeded (98 μL) into wells of a 96-well microtiter plate. Two-fold serial dilutions of antibiotic or chemical compound were added in a volume of 2 μL. Plates were incubated overnight at 37°C.

### GFP Reporter Plasmids and cloning

Cpx, Rcs, and RpoD GFP transcriptional reporters were constructed by amplifying the promoter regions of *cpxP, osmB*, and *rpoD* using the primers listed in Table S3. Amplicons were used in Gibson assembly reactions with pUA66 (44) to generate plasmids. A Strep II affinity tag was introduced to the C-terminus of LolA using a PCR site-directed insertion strategy. The V24C substitution was introduced by PCR site-directed mutagenesis.

### GFP Reporter Assays

Overnight cultures of GFP reporter strains were subcultured into fresh LB and 198 μL was seeded into black 96-well microtiter plates. Varying amounts of inducer or compound were added in a volume of 2 μL. Plates were grown at 37°C with shaking in a Synergy H1 plate reader (Biotek), and A_600nm_ and GFP fluorescence was measured every 10 mins. The amount of GFP per cell was calculated as a ratio of fluorescence to A_600nm_.

## Acknowledgements

This work was supported by grant 1R35GM133509 (to M.G.), fellowship F31AI147589 (to K.L.) and training grant T32AI106699 (to H.S.). We thank Daniel Wall (University of Wyoming), Nienke Buddelmeijer (Institut Pasteur), and Timothy Meredith (Pennsylvania State University) for providing LspA, Lgt, and Lnt depletion strains, respectively. We thank Benjamin Bratton (Vanderbilt University Medical Center) for providing *mreB* alleles. We are grateful to Kerrie May, William Shafer, and all members of the Grabowicz lab for helpful discussions and critical review of the manuscript. The authors declare no conflicts of interest.

## Supplemental Figure Legends

**Figure S1: Depletion of lipoprotein maturation or trafficking factors causes outer membrane permeability**. Strains in which the only copy of LspA, LolCDE, LolA, or LolB was under an arabinose inducible promoter were grown with increasing concentrations of inducer and the large scaffold antibiotics rifampicin and erythromycin. Depletion increased OM permeability, as measured by sensitivity to large scaffold antibiotics. Data are from three independent experiments; averaged density (A_600nm_) values of antibiotic-treated cultures relative to mock-treated control (set as 1.0) are shown.

**Figure S2: Deletion of *lpp* does not protect against general OM biogenesis defects**. Relative viability of strains with arabinose-dependent expression of *bamD* or *lptE*. Viable counts per mL of culture were enumerated in the presence of 0.2% arabinose or in the absence of arabinose and used to determine the fold change in viability when inducer is absent. Data represent three independent experiments and the mean.

**Figure S3: Depletion of lipoprotein maturation or trafficking activates the Rcs stress response but does not activate general OM stress responses**. Inducible LspA, LolCDE, LolA, or LolB strains carrying *P*_*osmB*_*-gfp, P*_*micA*_*-gfp*, or *P*_*rpoD*_*-gfp* were grown with (black solid) or without (blue dots) inducer. Culture density (OD_600_) and fluorescence were measured to calculate fluorescence per cell (Fluorescence/OD_600_). (Data are average ± standard deviation, N=3).

**Figure S4: Chemical inhibitors of OM lipoprotein biogenesis cause Rcs activation but do not activate *σ***^**E**^ **or RpoD**. Stress response activation after treatment (arrow) with DMSO (black), MAC13243 (orange), A22 (red), globomycin (blue), or compound 2 (purple) was assessed in a Δ*tolC* background. Culture density (OD_600_, top) and fluorescence were used to calculate fluorescence per cell (OD_600_/Fluorescence) in strains carrying *P*_*osmB*_*-gfp, P*_*micA*_*-gfp*, or *P*_*rpoD*_*-gfp*. (Data are average ± standard deviation, N=3).

**Figure S5: MTSES inhibits the activity of LolA(V24C) and resistance to MTSES increases in the absence of Lpp**. A) Growth (OD_600_) of strains carrying either *lolA*^*+*^ (black) or *lolA(V24C)* (blue) was measured upon treatment with MTSES. In early log phase (arrow), strains were treated with 0.5 mM MTSES (dotted) or vehicle control (1% DMSO) (solid). (Data are average ± standard deviation, N=3). B) MICs of strains carrying either *lolA+* or *lolA(V24C)* were assessed in the presence (*lpp*+) and absence (Δ*lpp*) of Lpp (N=3).

**Figure S6: An allele specific inhibitor of LolA causes OM permeability to antibiotics**. OM permeability to vancomycin, novobiocin, rifampicin, and erythromycin was assessed in *lolA*^*+*^ strains upon treatment with increasing amounts of MTSES (top). OM permeability to rifampicin and erythromycin were also assessed in a strain carrying *lolA(V24C)* upon increasing concentrations of MTSES (bottom). Data are from three independent experiments; averaged density (A_600nm_) values of antibiotic-treated cultures relative to mock-treated control (set as 1.0) are shown.

**Figure S7: MTSES treatment activates the Cpx stress response**. Cpx activation upon treatment with MTSES was assessed in a *lolA*^*+*^ background. Growth (OD_600_) of the *lolA*^*+*^ strains carrying a Cpx reporter plasmid (*P*_*cpxP*_*-gfp*) was measured. Strains were either *nlpE*^+^ (blue) or *nlpE::spec* (orange). In early log phase (arrow), strains were treated with DMSO (1%) (solid) or 0.5 mM MTSES (dotted). Fluorescence per cell was calculated using fluorescence normalized to OD_600_. Reported values are normalized to a DMSO-treated control (Data are average ± standard deviation, N=3).

**Figure S8: MTSES treatment causes activation of the Rcs stress response**. Cpx activation upon treatment with MTSES was assessed in *lolA*^*+*^ (blue) and *lolA(V24C)* (orange) strains carrying *P*_*osmB*_*-gfp, P*_*micA*_*-gfp*, or *P*_*rpoD*_*-gfp*. Growth (OD_600_) and fluorescence were measured after treatment with DMSO (1%) (solid) or 0.5 mM MTSES (dotted) in early log phase. Fluorescence per cell was calculated by normalizing fluorescence to OD_600_. Reported fluorescence values are normalized to a DMSO-treated control (Data are average ± standard deviation, N=3).

